# Sex Differences in Age-related Impairments Vary across Cognitive and Physical Assessments in Rats

**DOI:** 10.1101/718361

**Authors:** Abbi R. Hernandez, Leah M. Truckenbrod, Keila T. Campos, Sonora A. Williams, Sara N. Burke

**Author notes:** Indicates corresponding author.

## Abstract

Inclusion of female subjects in biomedical research is imperative for understanding the mechanisms of age-related cognitive decline, as more than half of individuals older than 65 are female. Few behavioral and physical assays, however, have been conducted in both sexes within the same study. In the current experiments young and aged male and female rats underwent a battery of cognitive and physical assessments to examine for potential sex and age differences. Physical performance was measured with a rotarod test of motor coordination, assessment of maximum grip strength and swim speed. While there were differences between males and females in rotarod and grip strength, there was also a clear age-dependent decline in physical performance in both sexes. Cognitive assessment included the Morris watermaze test of hippocampal dependent spatial navigation and a biconditional association task (BAT) with a working memory (WM) component. Notably, a similar BAT has previously been validated as a more sensitive assay of age-related cognitive decline than the watermaze in male rats, which is replicated here in both female and male rats. Furthermore, young and aged female rats both spent a similar percent of time in each estrus cycle phase and phase did not influence WM/BAT performance. In conclusion, robust age differences in biconditional association task performance are observed in both sexes with no need for increased cohort size. Thus, future studies utilizing similar behavioral paradigms should be representative of the human population they intend to model through the inclusion of female subjects.

## 1. Introduction

The importance of including females in clinical research gained a lot of attention in the early 1990s, resulting in a balancing of female and male participants in NIH-funded clinical research [1]. Unfortunately, a commensurate equality of females in basic and preclinical research did not accompany this change [2,3]. In 2014, the directors of the US National Institutes of Health and of the US National Institutes of Health Office of Research on Women’s Health published an article in the journal *Nature* calling for the inclusion of female subjects and consideration of sex as a biological variable [1]. This was formalized as an NIH-implemented policy, such that the consideration of sex as a biological variable became a reviewable criteria in grant proposals beginning in 2016 [4] (NIH NOT-OD-15-102).

Though there has been an increase in the number of studies utilizing females, there is still a dearth of information regarding how females differ from males on commonly utilized cognitive/behavioral tasks, such as object discriminations, as well as common measures of physiological health [2]. Further complicating this issue are the numerous perceived barriers against the inclusion of females. This includes increased cost associated with larger animal numbers, the erroneous perception of increased behavioral and physiological variability of females relative to males [5], as well as the lack of available females of certain rodent strains from common breeders, including the National Institute on Aging [6,7]. Despite these barriers, several key studies have documented differences in social behavior and behavioral outcomes, as well as the underlying molecular neurobiology across male and female rats (reviewed in [8]). Sex differences within the context of age-related cognitive and physical decline, however, remain largely under investigated. Evaluating potential behavioral differences between males and females across the lifespan is critically important, as females represent a larger proportion of older adults and have a longer mean life expectancy relative to males [9].

Although no overt differences between males and females in cognitive faculties have been documented across mammals [10], a reliable, but modest, performance advantage of males compared to females on spatial tasks has been reported in both rodents [11] and humans [12]. A widely used behavioral assay for testing spatial learning and memory across the lifespan in rodents is the hippocampus-dependent spatial version of the Morris watermaze task, in which animals learn the location of a hidden escape platform in a tank of cold water over several days [13–15]. Although a limited number of studies have directly compared intact male and female rodent performance on the Morris watermaze, the results have been equivocal, particularly in regards to the age at which deficits emerge [16–18]. While variations in rodent strain and methodology could explain these discrepancies [19], across studies it is clear that in both females and males, aged rodents perform worse on the spatial version of Morris watermaze compared to their younger counterparts.

Previously, we have reported that an object-place paired biconditional association task is particularly sensitive to age-related cognitive decline in male rats, detecting age differences earlier in the lifespan than the Morris watermaze [20]. This task requires rats to alternate through two different arms of a maze, making a distinct object discrimination within each arm for a small food reward. While the objects remain the same throughout testing, the correct choice varies as a function of location within the maze. As this task requires the integration of object information with spatial location to flexibly update behavior, it depends on connectivity between the prefrontal cortex, perirhinal cortex and hippocampus [21–23]. Female rats have been documented as performing differently from males on other prefrontal cortical dependent behaviors, such as deliberative decision making in the face of risk of punishment [24]. Thus, it is critical to evaluate female cognition across the lifespan on behaviors that are prefrontal cortical-dependent and that have been shown to decline in aged males.

When examining sex and age differences, one variable that could confound the interpretation of data is physical ability, as peripheral health correlates strongly with cognitive capabilities [25–28] and sensorimotor impairments could impact performance. In fact, decreased performance on physical tasks predicts poorer cognitive outcomes [29–31]. While declining physical health and motor deficits associated with advanced age could lead to behavioral performance declines independent of cognitive status, there has been little work done to compare motor function of male and female rats across the lifespan. The current study, therefore, characterized young and aged male and female rats on a cognitive and physical test battery. While a robust age effect was replicated on a variant of the object-place association task called the working memory/biconditional association task (WM/BAT), there was no performance difference between males and females. Furthermore, WM/BAT performance did not differ across different phases of the estrus cycle in female rats. In contrast, males outperformed females on the early training blocks of the Morris watermaze task in both age groups, but this sex difference was absent in the later training blocks. Moreover, retention of platform location assessed by probe trials did not vary between males and females or age group. Females did perform significantly better than males on a rotarod test of motor coordination at all ages, although there were still significant effects of age in both sexes. Importantly, the physical advantage of females relative to males on the rotarod test did not confer a sex difference on the cognitive assessments.

## 2. Materials and Methods

### 2.1 Subjects & Handling

Young (4-7 months) and aged (23-24 months) male (n = 10) and female (n = 10) Fisher 344 x Brown Norway F1 (FBN) Hybrid rats from the NIA colony at Charles River were used in this study (n = 5/group; 20 rats total). Each rat was housed individually and maintained on a 12-hr light/dark cycle and all behavioral testing was performed in the dark phase. To encourage appetitive behavior in discrimination experiments, rats were placed on restricted feeding in which 20 ± 5 g (1.9 kcal/g) moist chow was provided daily and drinking water was provided *ad libitum*. Shaping began once each rat reached approximately 85% of their weight that corresponded to an optimal body conditional score of 3 [32]. All procedures were in accordance with the NIH Guide for the Care and Use of Laboratory Animals and approved by the Institutional Animal Care and Use Committee at the University of Florida.

### 2.2 Physical Assessments

Prior to behavioral testing and restricted feeding, rats’ physical capabilities were assessed. Grip strength (kgF) was tested using a Chatillon Grip Strength apparatus (Columbus Instruments; Columbus, OH). Rats were held by the tail and allowed to place their forelimbs on the machine. Once the rat was able to grip the machine, they were firmly pulled away to measure grip strength. This was repeated for a total of three trials and the maximum value was recorded for each rat. Rats were weighed immediately prior to testing.

A subset of 6 young and 6 aged rats (half female) were trained on the rotarod test of motor coordination (Rotamex, Columbus Instruments; Columbus, OH) for 2 consecutive days at 4.0 RPM. On the 3rd and 4th days, motor performance was evaluated by an accelerating rotarod test consisting of 6 total trials across 2 days. The trial began at 4.0 RPM and accelerated 1.0 RPM every 8.0 seconds for a maximum of 300 seconds. The time spent on the rotarod (seconds) and the speed of rotation at the time of fall (RPM) were recorded. In addition to grip strength and motor coordination, the swim velocity during the visually cued trials of the watermaze were recorded as a final assessment of physical performance using the software Water 2100 (HVS Image).

### 2.3 Morris Watermaze Test of Spatial Reference Memory

Spatial learning and memory performance was tested with the Morris watermaze as previously described [20,33]. Briefly, each rat received 3 trials per day, at a maximum of 90 seconds per trial, for 8 consecutive days. During each trial, the platform was hidden 2 cm below the surface of opaque water. Probe trials, in which the platform was removed for the first 30 seconds of the trial, were conducted during the last trial of the day on every other day to assess the proximity of the swim path to the target location (cumulative search error; see [34]). Performance was divided into 4 training blocks, with each block consisting of the 5 trials between each probe trial. Training blocks were used to calculate a Spatial Learning Index (SLI), which is a weighted sum of the cumulative search error during probe trials 2–4 (Spatial Learning Index = probe 2*1.25 + probe 3*1.6 + probe 4*1.7) as previously described [34]. At the end of block 4, a final test day consisting of 6 trials in which the platform was visible was used to ensure that no rats had impaired sensorimotor function. Swim speed during these trials was also used to assess physical performance (see section 2.2 above).

### 2.4 Apparatus and Habituation

Alternation training and working memory/biconditional association task (WM/BAT) behavioral testing occurred on a figure 8-shaped maze (see Figure 1) that was 67.5 inches long and 25 inches wide. The maze was constructed from wood and sealed with waterproof white paint. The center arm was made of clear acrylic. The choice platforms each contained two food wells (2.5 cm in diameter) that were recessed into the maze floor by 1 cm. All arms were 4 inches wide. The choice platforms were contained within 7.5 cm raised walls and the right arm was contained within 20 cm high raised walls, but the center and left arms did not have walls. Thus, the arms of the maze had an asymmetry, such that only the right arm was enclosed while the animals were relatively more exposed on the middle and left arms. This asymmetry biases rats to alternate towards the closed ‘safe’ arm [35]. To dampen the influence of extraneous noise on behavior, a white noise machine was used during behavioral training and testing. Rats were habituated to the testing apparatus for 10 minutes a day for 2 consecutive days, with Froot Loop pieces (Kellogg Company, Battle Creek, MI) scattered throughout the maze to encourage exploration.

**Figure 1:**
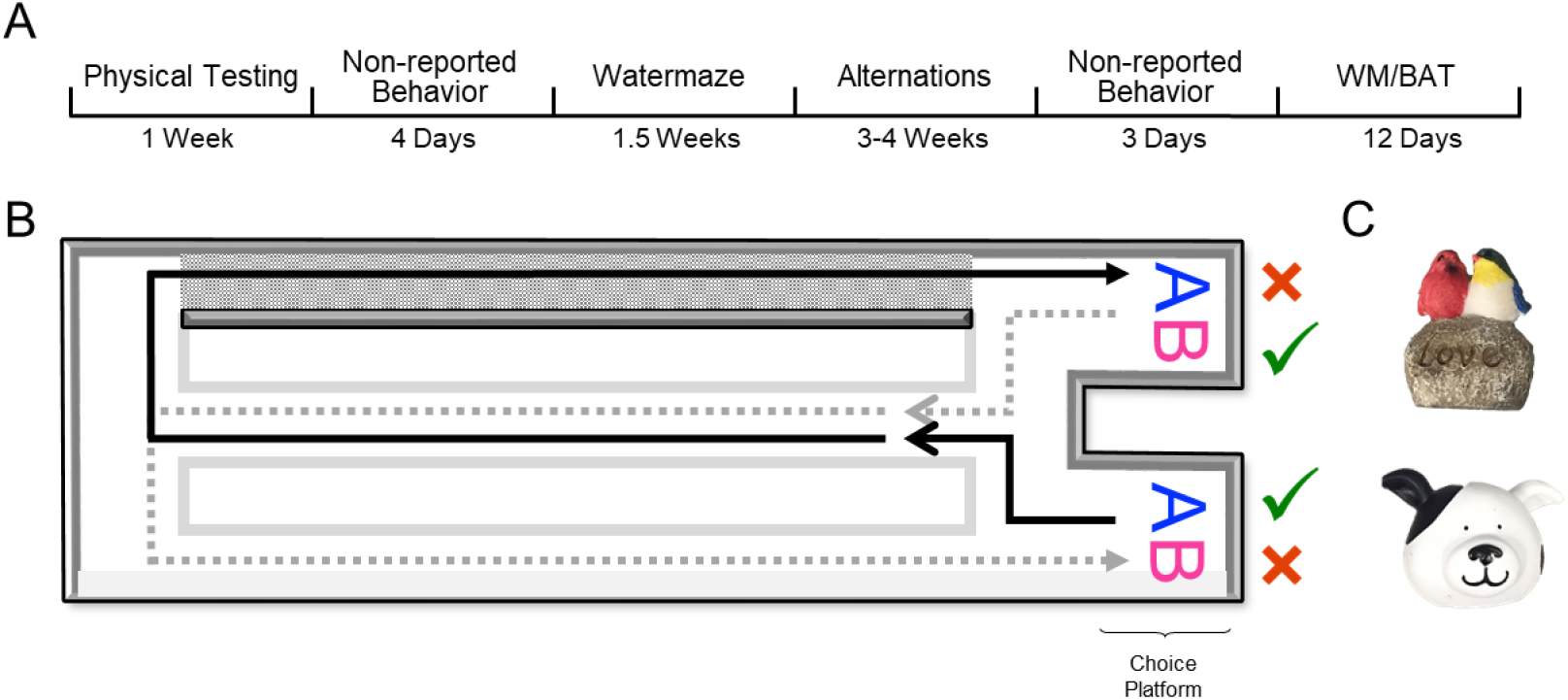
Behavioral testing paradigm. A) Experimental timeline. Note that spontaneous object recognition behavioral paradigms are not reported within this manuscript and these tasks occurred prior to Watermaze and WM/BAT testing. B) A figure-8 shaped maze was used for alternation training and the working memory/biconditional association task (WM/BAT; birds eye view of the maze). In this task, rats had to alternate between trials in the left and right arms of the maze, with the correct object choice contingent upon location (i.e. object A is correct within the left arm, object B is correct within the right arm) regardless of which food well the object is covering (left or right food well within the choice platform). Gray area indicates closed ‘safe’ arm of the maze (rightward turn). C) Objects used for WM/BAT task.

### 2.5 Working Memory/Biconditional Association Task (WM/BAT)

Following habituation to the testing apparatus, rats were trained to alternate between the left and right arms of the maze. Correct alternations were rewarded with ~1/2 of a Froot Loop placed randomly in either well within the choice platform. When rats were alternating correctly ≥80% of the time on 2 consecutive days, they began training on an object discrimination-based task. On the first day of training with objects, the objects only partially covered the food wells on the first 8 trials (4 per arm) to facilitate the rats learning that food was hidden beneath objects. In between alternation training and WM/BAT testing, rats performed a simple (non-biconditional) object discrimination task not reported here.

Rats were trained on the working memory/biconditional association task (WM/BAT) as previously described [35] for 12 consecutive days using 2 novel objects. In addition to correctly alternating between the left and right arms of the maze, rats had to choose between a pair of objects presented in the choice platform. The same object pair was presented in both arms, and the correct object choice was contingent upon the location within the maze. For example, rats presented with objects A and B must choose object A in the left arm and object B in the right arm to receive a Froot Loop reward. For all days of WM/BAT training, each testing session consisted of 20 trials per day and incorrect alternations resulted in removal of objects (no object choice presented to the rat) until the next turn on which they correctly alternated throughout the maze. If a rat alternated incorrectly 3 times in a row, they were guided to the correct side on the subsequent turn to ensure objects were encountered on an adequate number of trials per training session.

### 2.6 Estrus monitoring

To investigate whether the female rats’ performance was affected by their menstrual phase, all rats were retested on the WM/BAT following the completion of the behavioral testing described above and the estrus cycle phase was determined each day following behavioral testing. Throughout object discrimination testing, female estrus cycle was monitored. Following completion of behavioral testing each day, a small amount of sterile saline was used to produce vaginal smears placed on glass slides. Unstained slides were examined at 40X using light microscopy, and estrus cycle phase was determined using criteria outlined in [24,36].

### 2.7 Statistical analyses

All data are expressed as group means ± standard error of the mean (SEM) unless otherwise reported. Morris watermaze spatial learning index, path length and latency, rotarod latency and speed, grip strength, percent of correct object/direction choices for each task and number of incorrect trials were analyzed using a two-factor ANOVA with the between-subjects factors of age (2 levels: young and aged) and sex (2 levels: male and female). Tasks in which the path length, the percent of correct object choices or correct alternation choices were compared across multiple days or multiple arms were analyzed using repeated measures-ANOVAs (RM-ANOVA) with the between-subjects factors of sex and age. Estrus cycle effects on performance were analyzed via RM-ANOVA across each phase of the cycle with the between-subjects factor of age. Finally, to examine for potential relationships across variables, a principal component analysis (PCA) was performed with a varimax rotation. Factors with eigenvalues above were considered meaningful and loading coefficients below 0.40 were excluded. The null hypothesis was rejected in all cases when p-values were < 0.05. All analyses were performed with Statistical Package for the Social Sciences (SPSS) v25 or GraphPad Prism version 7.03 for Windows (GraphPad Software, La Jolla, California USA).

## 3. Results

### 3.1 Aged-related declines in physical performance

To assess physical performance decline, and the extent to which this could interact with cognitive outcomes, 3 rats in each sex and age group first completed a rotarod test of sensorimotor impairment, and grip strength was evaluated in all rats. Univariate ANOVA for the latency to fall off of the rotamex machine that was used to test rotarod performance indicated that aged rats fell off in significantly less time than young rats (F_[1,8]_ = 28.31; p < 0.001) and that male rats fell off in significantly less time than female rats (F_[1,8]_ = 52.48; p < 0.001; Figure 2A). There was a trend for an interaction between age and sex that did not reach significance (F_[1,8]_ = 4.24; p = 0.07), though a previous report has shown aged female mice aren’t as susceptible to age-related performance deficits at rotarod tasks as their male counterparts [37], which is consistent with the observations reported here. Univariate ANOVA for the speed at which rats fell off of the rotarod indicated aged rats fell off at significantly slower speeds than young rats (F_[1,8]_ = 35.01; p < 0.001) and that males fell off at significantly slower speeds than females (F_[1,8]_ = 65.34; p < 0.001; Figure 2B). There was no interaction between age and sex, but a trend towards an effect (F_[1,8]_ = 3.68; p = 0.09).

**Figure 2:**
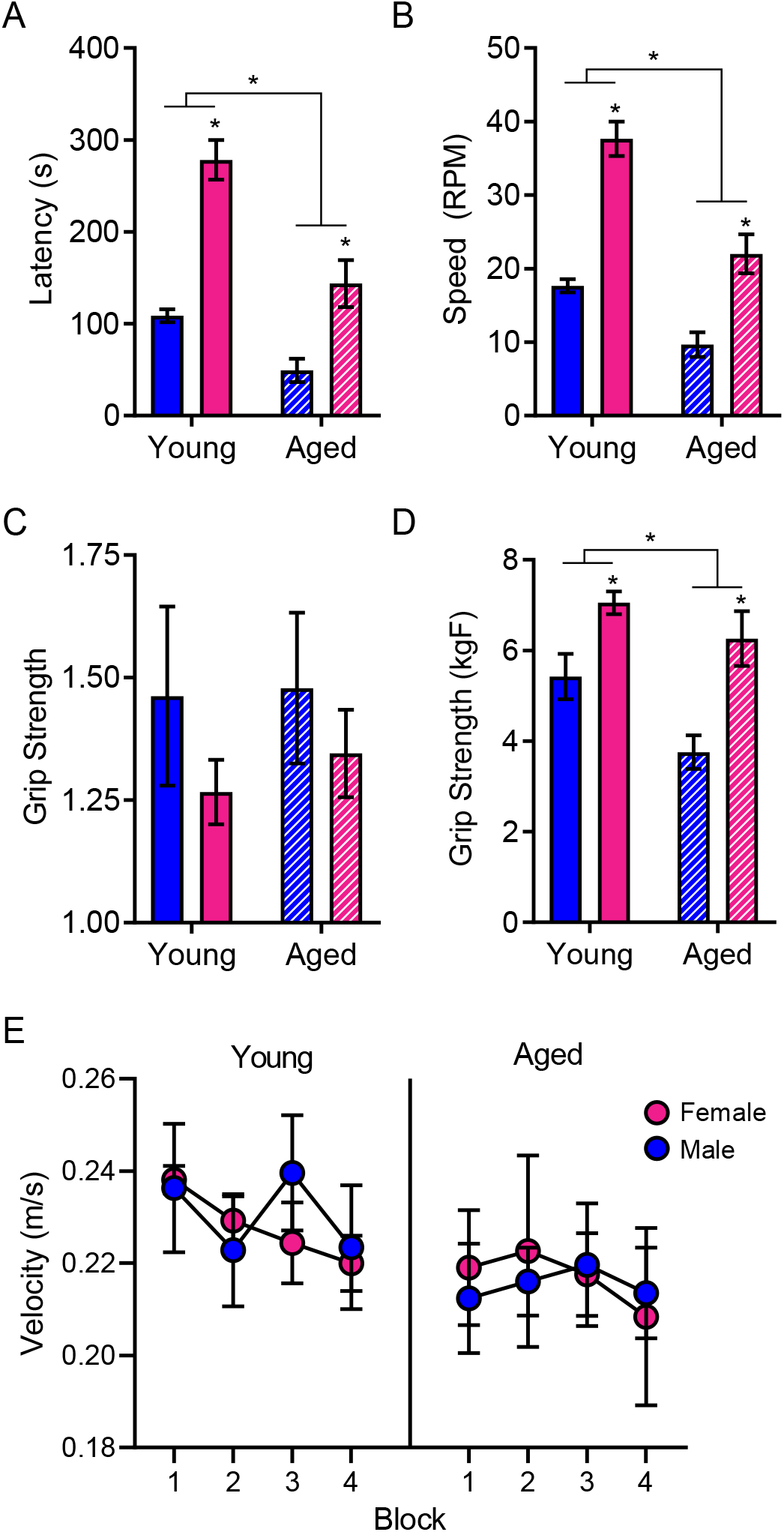
Aged rats demonstrated physical impairments relative to young rats. A) Rotarod testing demonstrated that aged rats fell off in significantly less time than young rats and female rats stayed on significantly longer than males. B) Furthermore, aged rats fell off at a significantly slower speed than young rats and female rats fell off at significantly higher speeds than males. C) While there were no differences in raw grip strength across groups, D) when normalized to body weight, females outperformed males and age significantly impaired grip strength in both sexes. E) There were no differences across groups on swim speed during the Morris watermaze. All data represents group means ± SEM.

Raw grip strength did not differ by age (F_[1,16]_ = 0.13; p = 0.73) or sex (F_[1,16]_ = 1.56; p = 0.23), and there was no significant interaction between age and sex (F_[1,16]_ = 0.06; p = 0.82; Figure 2C). Grip strength, when normalized for body weight, was significantly greater in females than it was for males (F_[1,16]_ = 20.91; p < 0.001) and significantly declined with age (F_[1,16]_ = 7.40; p = 0.02). This is due to the smaller size of female rats, and the increased mass of aged males. There was no significant interaction between age and sex (F_[1,16]_ = 0.94; p = 0.35), demonstrating grip strength was affected by aging in a similar manner across both sexes.

In addition to the rotarod and grip strength measures, the Morris watermaze can be used to assess physical performance by quantifying the swimming velocity on the cued trials in which the escape platform is visible and the rat is swimming towards an obvious goal with little uncertainty. RM-ANOVA on blocks 1-4 with the between-subjects factors of age and sex revealed no significant main effect of training block (F_[3,48]_ = 1.90; p = 0.14), age (F_[1,16]_ = 1.51; p = 0.24) or sex (F_[1,16]_ < 0.01; p = 0.96) on swim velocity, nor were there any significant interactions between these variables (Figure 2D).

### 3.2 No behavioral differences were detected on the hippocampal-dependent Morris watermaze by age or sex

Rats were tested on the Morris watermaze as described in previous experiments [20,33,34,38] to determine whether age or sex were related to hippocampal-dependent spatial memory performance. Previously, we have observed that male rats of this age and strain do not show robust age-associated impairments on this protocol of the Morris watermaze task between 4 and 24 months. The corrected integrated path length (CIPL; also known as cumulative search error) accounts for differences in velocity across subjects and measures the swim path to the platform compared to the most efficient path. RM-ANOVA on CIPL from blocks 1-4 with the between-subjects factors of age and sex revealed a significant main effect of training block (F_[3,48]_ = 12.23; p < 0.001), which trended towards significantly interacting with age (F_[3,48]_ = 2.75; p = 0.053). Training block did not significantly interact with sex (F_[3,48]_ = 0.72; p = 0.55; Figure 3A), suggesting that both males and females improved across training. The main effect of sex on Morris watermaze performance did reach statistical significance (F_[1,16]_ = 6.87; p = 0.02), but there was no significant effect of age (F_[1,16]_ = 2.63; p = 0.13). Moreover, the interaction between age and sex did not reach statistical significance (F_[1,16]_ = 1.06; p = 0.32). Notably, the significantly decreasing path length across training blocks for all groups indicated that rats in all groups were able to learn the platform location (see table 1 for comparisons). The degree to which the path length decreased from block 1 to block 4 did not significantly differ by age (F_[1,16]_ = 1.17; p = 0.30) or sex (F_[1,16]_ = 0.80; p = 0.39), nor did these 2 factors interact significantly (F_[1,16]_ < 0.001; p > 0.99). These data suggest that the sex differences in Morris watermaze performance were most evident in the early training blocks and that by the 3^rd^ and 4^th^ training blocks, the differences between males and females of both age groups were diminished.

**Table 1:**
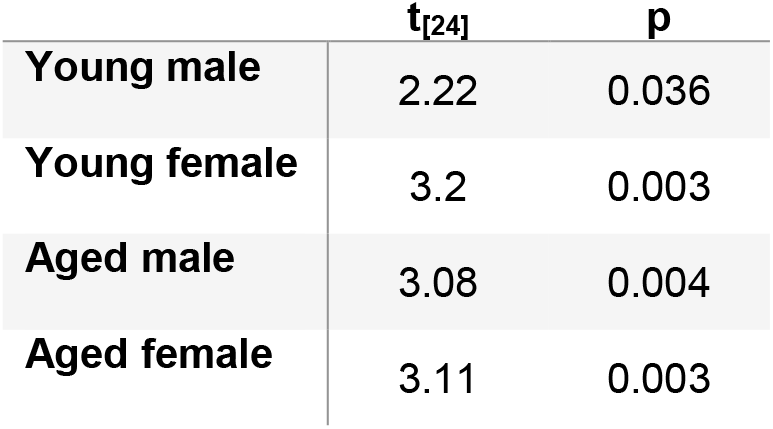
Paired samples t-tests on corrected integrated path length (CIPL) between blocks 1 and 4 for each group indicated significantly decreasing path lengths for all groups throughout training.

| | $t_{[24]}$ | $p$ |
| --- | --- | --- |
| **Young male** | 2.22 | 0.036 |
| **Young female** | 3.2 | 0.003 |
| **Aged male** | 3.08 | 0.004 |
| **Aged female** | 3.11 | 0.003 |

**Figure 3:**
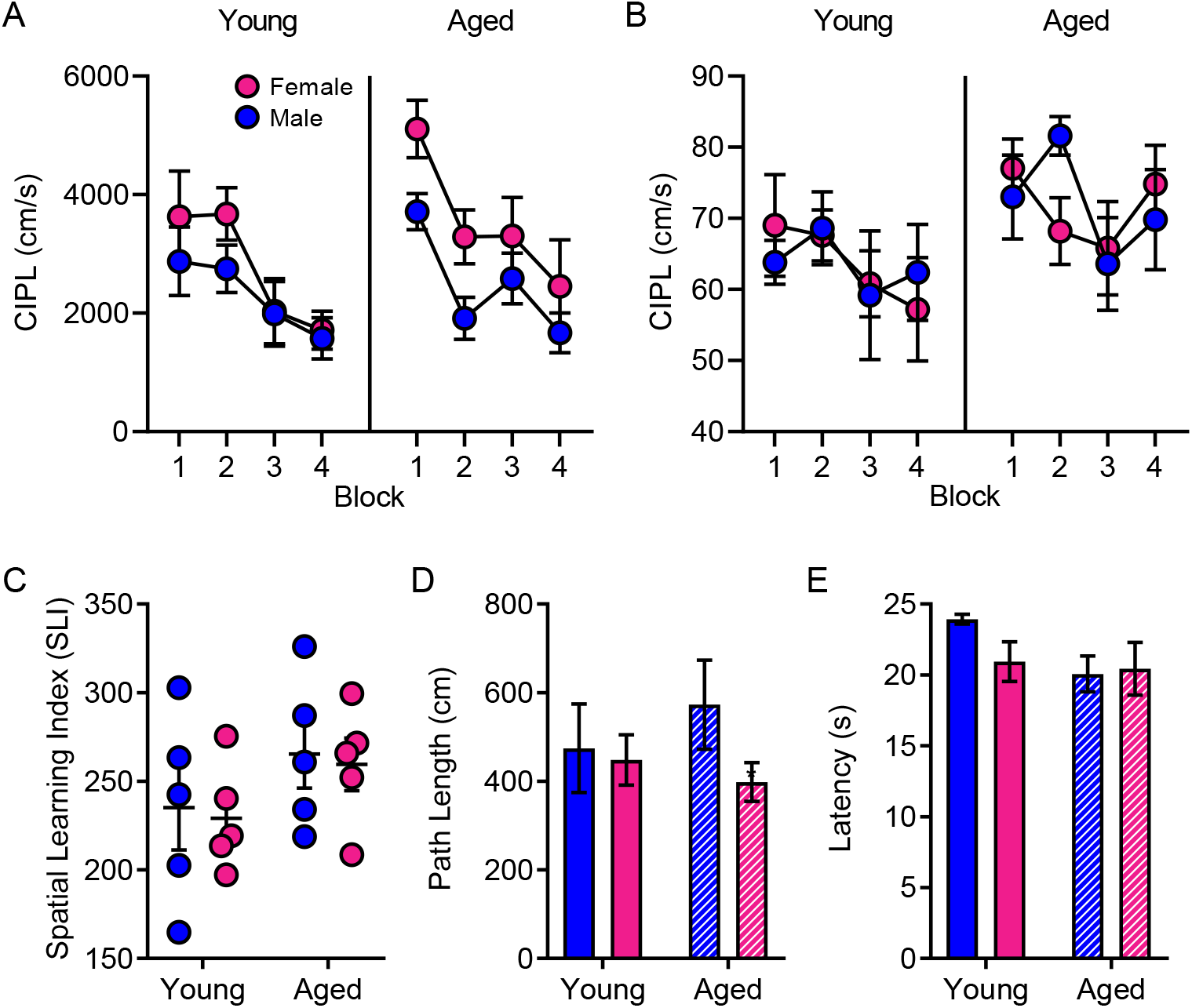
Morris watermaze performance by age and sex. A) The corrected integrated path length (CIPL) during training trials did not differ across age or sex. B) Furthermore, the CIPL values during probe trials and C) the spatial learning index (SLI) were also unaffected by either age or sex. D) During cue training, there were no differences in path length or E) latency to find the platform across groups. Data in parts A, B, D & E represent group means ± SEM.

CIPL was also calculated for each probe trial (1 per block). RM-ANOVA on the CIPL for all 4 probe trials with the between-subjects factors of age and sex revealed a significant main effect of trial (F_[3,48]_ = 3.44; p = 0.02). Probe trial did not significantly interact with age or sex (F_[3,48]_ = 1.20; p = 0.32), nor was there a significant main effect of sex (F_[1,16]_ < 0.01; p = 0.96) on probe trial performance. The main effect of age did, however, trend towards significance (F_[1,16]_ = 3.62; p = 0.08). Consistent with the lack of an age effect on training and probes trials, the Spatial Learning Index (SLI) was also not significantly different between young and aged rats (F_[1,16]_ = 2.73; p = 0.12) or between sexes (F_[1,16]_ = 0.11; p = 0.75; Figure 3C). Finally, there was no significant interaction between age and sex on SLI values (F_[1,16]_ < .001; p > 0.99). Together these data indicate the aged rats utilized in this study were not impaired on the hippocampus-dependent Morris watermaze task, which is consistent with previous reports in this rat strain [20].

Cue trials in which the platform was visible to the rats were performed to exclude the possibility of sensorimotor impairments. There were no differences by age (F_[1,16]_ = 0.09; p = 0.76) or sex (F_[1,16]_ = 1.61; p = 0.22) on the average path length during cue training (Figure 3D). Furthermore, there were no differences between age (F_[1,16]_ = 2.69; p = 0.12) or sex (F_[1,16]_ = 0.98; p = 0.34) on the average time it took to find the platform during cue training (Figure 3E). There were no significant age by sex interactions for path length or latency (p > 0.22 for both comparisons).

### 3.3 Behavioral shaping was impaired by age but was not affected by sex

Prior to the introduction of objects on the working memory/bi-conditional association task (WM/BAT), rats were first trained to alternate between the left and right arms of the maze. Because learning effects may be masked by procedural acquisition or differences in the amount of time it took aged rats to become appetitively motivated, the first 2 days of alternation training were excluded from analysis, although results were similar when these days were included. RM-ANOVA on days 3-10 of testing revealed a significant main effect of day (F_[7,112]_ = 2.41; p = 0.02), but no interactions between day and age or sex (p > 0.67 for both comparisons). While there was no main effect of sex (F_[1,16]_ = 2.87; p = 0.11), there was a significant main effect of age (F_[1,16]_ = 5.65; p = 0.03), such that young rats acquired the alternation rule more quickly than the aged animals.

During alternation training, because of the asymmetry of the maze, all rats had a predisposition against turning in the leftward direction towards the open arm, rather than alternating evenly between the two arms. The tendency to turn right over left can be used to calculate a turning bias. Across days of testing, RM-ANOVA indicated a main effect of day on the turn bias (F_[22,352]_ = 2.49; p < 0.0001), but neither age (F_[1,16]_ = 1.73; p = 0. 21) nor sex (F_[1,16]_ = 2.66; p = 0.12) significantly affected the turn bias, and there was not a significant interaction between age and sex on the turn bias (F_[1,16]_ = 1.83; p = 0.20). These data show that males and females of both age groups were similarly predisposed to traverse the closed over the open arm.

### 3.4 Working memory/biconditional association task (WM/BAT) performance is impaired by age but does not differ by sex

Rats were also trained on a cognitive multi-task that required both an object discrimination and an association of an object with a particular location within the maze. This type of biconditional association task is particularly sensitive to age-related cognitive decline [20]. The object-place paired association task was combined with a continuous spatial alternation test of working memory to increase the cognitive load. Combining these two behaviors into one task results in an assessment of cognitive multi-tasking referred to as working memory/biconditional association task (WM/BAT). Learning effects on this task may be masked by procedural acquisition during the first few days of training. Moreover, during the first few days of training the food reward was made visible to facilitate the rats learning to push objects to retrieve the food reward from the well underneath. Thus, the first 2 days of WM/BAT acquisition/procedural training were not included in the analysis. RM-ANOVA on days 3-12 of testing revealed a significant main effect of day (F_[9,144]_ = 2.01; p = 0.04), but no interactions between day and age or sex (Figure 4A). Similar to previously published results, aged rats performed significantly worse than young rats at this task (F_[1,16]_ = 8.66; p = 0.01). Male and female rats, however, did not significantly differ (F_[1,16]_ = 0.07; p = 0.80), nor was there an interaction between age and sex (F_[1,16]_ = 0.10; p = 0.76). Moreover, the number of incorrect trials on days 3-12 of training were tabulated, and young rats had significantly fewer incorrect trials than aged rats (F_[1,15]_ = 7.55; p < 0.02). Importantly, there was no difference between male and female rats on the number of incorrect trials (F_[1,15]_ = 0.81; p = 0.38), nor did age and sex significantly interact (F_[1,15]_ =0.11; p = 0.74) on this performance measure.

**Figure 4:**
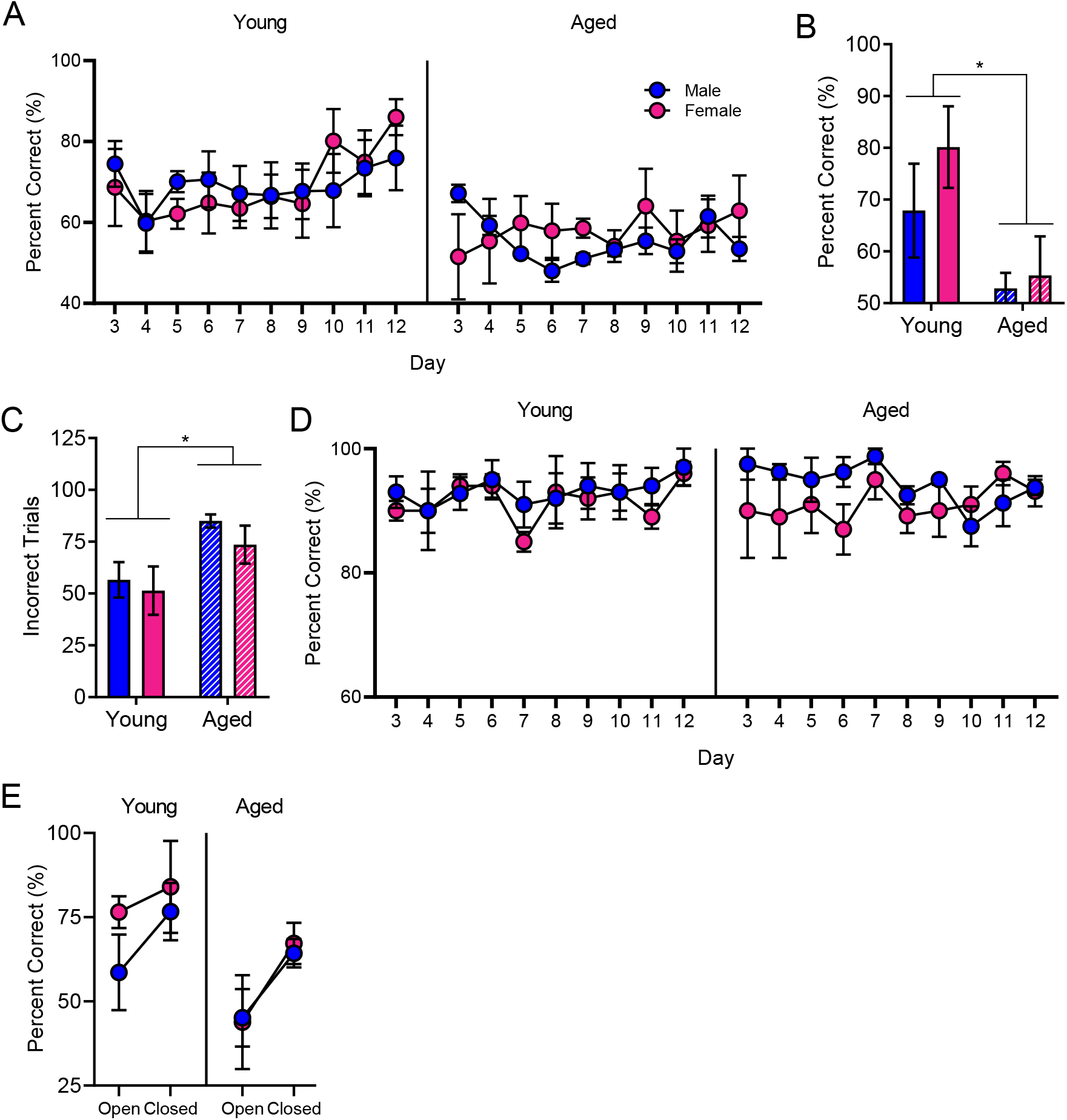
Working memory/biconditional association task (WM/BAT) performance. A) Aged rats were impaired at WM/BAT acquisition relative to young rats. B) By the 10th day of testing, young rats of both sexes were performing significantly better than aged rats. C) Young rats performed significantly fewer incorrect trials overall across days 3-12 of WM/BAT training. D) Rats in all groups were alternating efficiently throughout WM/BAT testing, with no presentation of a side bias. E) Performance for trials within the closed arm was significantly better relative to performance during trials in the open arm for all groups. Data represent group means ± SEM.

Day 10 of testing was the first day that any group of rats reached a criterion performance of >80%. Thus, performance on this day was compared across groups. There was a significant effect of age (F_[1,16]_ = 7.56; p = 0.01), but not sex (F_[1,16]_ = 1.04; p = 0.32) on percent correct on day 10 of testing. Furthermore, the interaction between age and sex did not reach statistical significance (F_[1,16]_ = 0.46; p = 0.51), indicating that young rats were performing significantly better on this day of testing than aged rats regardless of sex (Figure 4B).

Lastly, the number of correct turns during WM/BAT were analyzed to assess working memory errors. RM-ANOVA across days 3-12 of WM/BAT testing indicated there were no differences with age (F_[1,15]_ =0.04; p = 0.84) or sex (F_[1,15]_ =1.87; p = 0.19) in the percent of correct alternations (Figure 4D). The lack of a significant effect of testing day (F_[1,15]_ =0.11; p = 0.91) on working memory errors, together with the fact there were no effects of day, sex or age on side bias during WM/BAT (p > 0.13 for all comparisons; data not shown), together indicate rats had learned to alternate correctly prior to beginning the WM/BAT task and did not get better or worse throughout subsequent days of training. Furthermore, this demonstrates that the added cognitive load of the WM/BAT task did not impair rats’ ability to alternate throughout the maze.

### 3.5 Anxiety-related cognitive deficit impaired rats of both ages and sexes

Due to the asymmetry of the maze, the effect of the anxiety-inducing open arm on cognitive performance, relative to performance within the closed arm, can be assessed. Because day 10 of WM/BAT training was the first day on which any group of rats reached a criterion performance of >80% correct, performances across the open and closed arms were compared on this day. RM-ANOVA with the repeated variable of arm of the maze (open versus closed) and between-subjects variables of age and sex revealed that all rats performed significantly worse in the open arm than they did in the closed arm (F_[1,15]_ = 7.40; p = 0.02; Figure 4E). While there was again a significant main effect of age (F_[1,15]_ = 6.01; p = 0.03), neither age (F_[1,15]_ = 0.06; p = 0.80) or sex (F_[1,15]_ = 0.46; p = 0.51) significantly interacted with arm, indicating that rats of all age groups and sexes experienced an anxiety-related cognitive deficit.

### 3.6 Behavioral performance was not related to estrus cycle phase

To investigate whether the female rats’ performance was affected by their menstrual phase, all rats were retested on the WM/BAT following the completion of the behavioral testing described above, and the estrus cycle phase was determined each day following behavioral testing (Figure 5). There was no effect of age on the length of time spent in each phase of the cycle (F_[1,25]_ = 1.65; p = 0.21; Figure 5A) nor on the duration of each cycle (t_[127]_ = 0.48; p = 0.64; Figure 5B), indicating that the aged females were still cycling. Because all behavioral training occurred during the rats’ dark phase, when rats are naturally active, the estrus phase of the cycle was not commonly observed (estrus was observed at least once in all but 1 young and 2 aged rats). Therefore, comparisons of behavioral performance were made across proestrus, metestrus and diestrus only. RM-ANOVA on the average performance during WM/BAT during each of the 3 cycle phases in female rats, with age as a between-subjects factor, indicated there was no significant effect of cycle phase on behavioral outcome (F_[2,16]_ = 1.79; p =0.20; Figure 5C). Furthermore, there was no significant effect of age (F_[1,8]_ = 0.57; p = 0.47), nor significant interaction between behavioral performance across each testing phase with age (F_[2,16]_ = 0.46; p =0.64).

**Figure 5:**
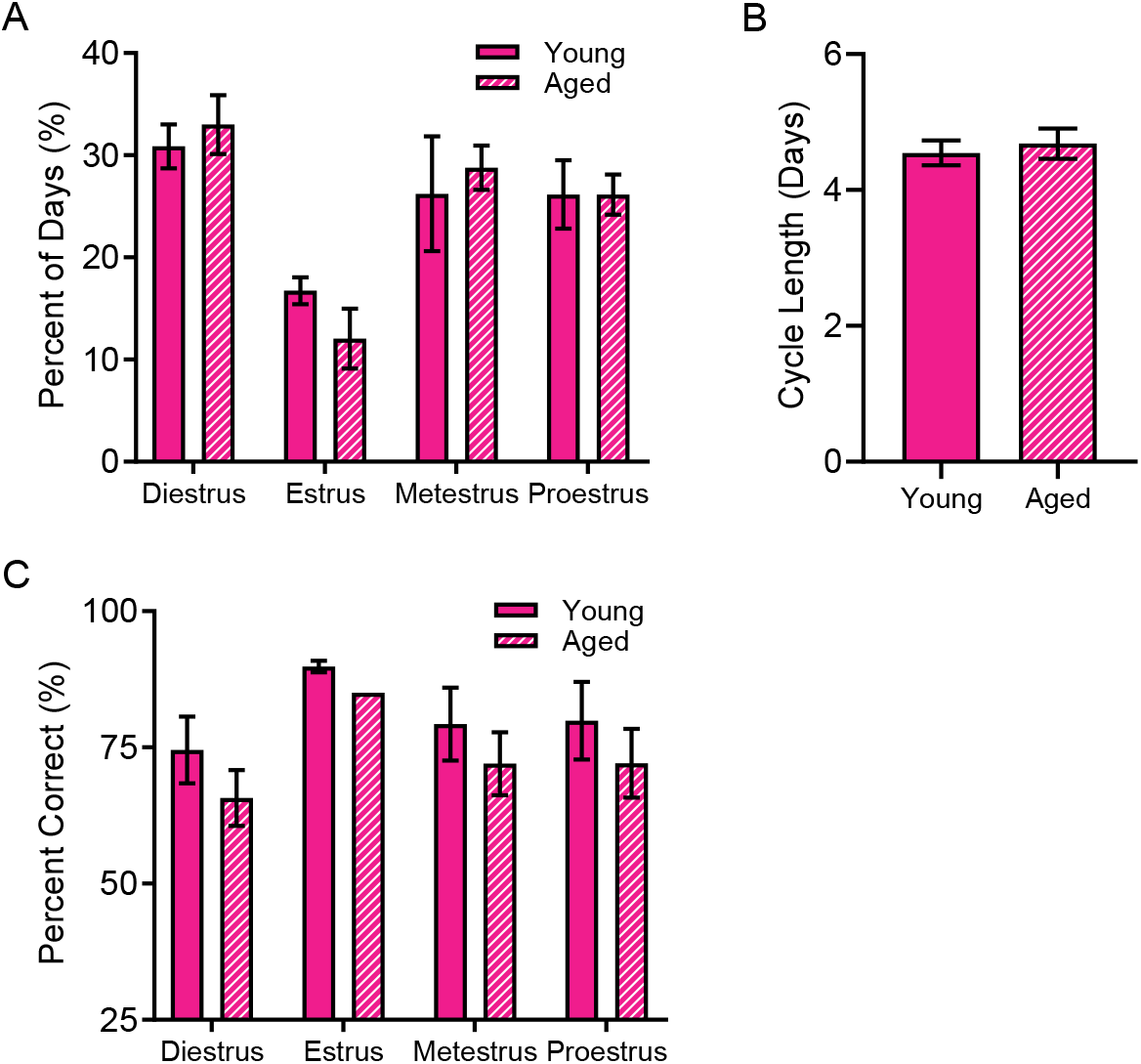
Estrus cycle interactions with age and behavioral performance. A) Age did not alter the percent of time that rats spent within each phase of the estrus cycle. B) Similarly, there were no alterations to the average cycle length across young and aged rats. C) Finally, the estrus cycle phase did not significantly interact with behavioral outcomes on the WM/BAT task. Data represent group means ± SEM.

### 3.7 Cognitive and physical performances are inversely correlated

A principle component analysis (PCA) was used to investigate a potential relationship between measures of cognitive and physical performance. The PCA, run with a varimax rotation to reduce split loadings, included spatial learning index (SLI) from the Morris watermaze, swim speed during cue trials of the Morris watermaze, incorrect trials to criterion during acquisition of alternations throughout the maze, incorrect trials on days 3-12 of WM/BAT testing, raw grip strength, weight at the time of grip strength testing and latency to fall off of the rotarod. There were no problematic redundancies for any of the variables (factor loadings > 0.22 across multiple components).

A model with 3 components explained 87.04% of the variance. The first component (eigenvalue = 3.74), which negatively loaded with both rotarod latency (−0.95) and grip strength (−0.88), accounted for 53.43% of the variance. Poorer physical performance on these tasks also strongly loaded positively with the number of incorrect trials before alternating correctly (0.94), indicating that rats who performed the worst on the rotarod task had the weakest grip strength, and were also the poorest at alternation task acquisition. Component 1 also loaded positively onto weight (0.88), indicating that larger rats were poorer performers on physical and cognitive tasks. The second component (eigenvalue = 1.30), which corresponded with incorrect trials to criterion on the WM/BAT task (0.79) and the SLI (0.71), accounted for 18.50% of the variance. Similar loading of these two components would be expected, as each of these tasks are dependent on the hippocampus. The third component (eigenvalue = 1.06) corresponded with swim speed during cue trials of the Morris watermaze (0.97) and accounted for 15.10% of the variance. No other variables loaded onto this component.

## 4. Discussion

In this study, we have replicated previous data showing that biconditional association tasks are particularly sensitive behavioral assays for detecting age-related cognitive decline relative to the Morris watermaze [20,35]. Furthermore, we observed that both males and females showed similar performance declines across age on the working memory/bi-conditional association task (WM/BAT) variant of this behavioral test. The current observation of no differences between males and females on WM/BAT performance validate and reinforce the necessity of futures studies incorporating female subjects into neurobiological investigations of the mechanisms of cognitive aging. Similarly, we found no differences in spatial memory recall during Morris watermaze testing and, as previously reported in rats of this strain and age, no age differences either [20]. While swim speed during cued trials of the Morris watermaze did not vary by sex, rotarod and grip strength did show sex-dependent differences in performance.

Although females comprise approximately 51% of the population in the United States [9], the inclusion of female subjects in studies utilizing animal models is not currently commonplace. In fact, there have been several arguments against the inclusion of females in biomedical and neuroscience research. Most notably, there is resistance against larger cohort sizes and increased cost of maintaining and testing larger colonies. However, we report here that including both young and aged female rats did not introduce increased variability due to sex differences and thus did not require an increased sample size relative to a previous study utilizing the same task with all male subjects [20]. There was no correlation with estrus cycle phase and behavioral performance, demonstrating that hormonal fluctuations throughout the cycle did not affect the cognitive outcomes measured here. In line with these data, there are several reports demonstrating that there is commonly no requirement for increasing sample size to incorporate females and furthermore no need to account for estrus cycle phase unless that is the focus of the study [39–42].

While there aren’t overt differences in cognitive level, there are slight differences across sexes observed in humans [43,44], non-human primates [45–47], and rats [11] that may need to be accounted for in foundational studies of cognitive aging in both sexes (for review see [48]). One such potential difference to take into account is anxiety, which has been reported to affect females more than males [49,50]. To assess this in our study, the figure-8 shaped maze (see Figure A) utilized for our behavioral paradigm was asymmetrically shaped such that the left arm of the maze had walls enclosing both sides of the arm (closed ‘safe’ arm) whereas the left arm was wall-less (open ‘risky’ arm), biasing rats to alternate rightward throughout the maze. While all rats were right arm-biased early in shaping, age was the only variable affecting alternation behavior and there was no difference across male and female willingness to enter the open arm. Furthermore, we did not observe a difference in working memory errors or open arm-induced cognitive deficit across sexes during WM/BAT, as both sexes and age groups were negatively affected by the open arm equally. The lack of effect during WM/BAT testing may be due to extensive shaping prior to starting the object choice tasks, potentially masking any sex-dependent differences in anxiety caused by the openness of the arm.

A second cognitive aspect that may differ between males and females is spatial learning and memory. Although the data presented here demonstrate no effect of sex on spatial memory retrieval during Morris watermaze performance in either age group, data from other studies have been equivocal regarding sex differences [16–18]. There are several possible explanations as to why data are inconsistent regarding sex differences in spatial navigation. Firstly, there may be differences in acquisition of the procedural aspects of the Morris watermaze, but no differences in spatial recall, as males typically outperform females at Morris watermaze acquisition, but don’t often demonstrate greater memory for location than females once the platform has been identified by the subject [11,48,51,52]. The current data are consistent with these previous observations, as females did not have impaired spatial learning indices, but had less efficient paths to the platform compared to males early in training. Secondly, the watermaze is a highly stressful task, as subjects are placed a novel environment from which they must swim to escape, often through cold water, and males and females may show different rates of habituation to stress. In support of this idea, factors reducing stress level also decrease the male advantage in multiple spatial navigation based tasks [53,54]. Furthermore, pretraining on Morris watermaze also ameliorates sex differences, further supporting a complex interaction between sex, stress and spatial learning [55].

Although hormonal influence on behavioral outcomes is often cited as a reason to exclude female subjects, as if hormones were a female deviation from male default state devoid of hormones, it is also true that male subjects undergo hormonal fluctuations that can affect behavioral outcomes. In species that are seasonal breeders, such as deer mice and meadow voles, the timing of Morris watermaze testing relative to the mating season influences the presence or absence of a male spatial advantage [51,52,56]. Furthermore, testosterone levels in males are highly dynamic and show greater variability in response to stress than female endocrine changes under similar conditions [57,58].

Finally, because many animal species frequently tested on spatial navigation ability via the Morris watermaze are nocturnal species, effects of circadian rhythm may influence the behavior of both sexes, as rodents behave differently during the light or dark phases of the day [59]. Several investigations showing robust sex differences on Morris watermaze performance were conducted during the light cycle [16], when rodents are normally sleeping, disrupting circadian rhythmicity. Other studies in which testing occurred during rodents’ dark phase, including the current data, demonstrate no differences between male and female performances [19]. These differences in time of testing may also influence stress levels particularly in aged subjects, as older mice demonstrate increased anxiety on light/dark testing [60].

In addition to assessing cognitive decline with age, physical strength and motor ability were also analyzed in these subjects. On a rotarod test of motor ability, young rats were able to maintain a higher speed and longer duration than aged rats, as previously reported [61–63]. Additionally, females strongly outperformed males in both age groups, demonstrating a clear sex difference in performance on this task. However, grip strength was lower in females relative to males when normalized for body size, suggesting the difference in rotarod performance was not due to heightened physical strength in the females, but rather could be related to differences in motor coordination. An alternative explanation for the sex difference in performance on the rotarod task is a discrepancy in motivation or valuation of risk of fall in males versus females. Female rats are more risk averse than males on a risky decision making task in which the choice of a larger food reward is associated with a mild foot shock, demonstrating they may be more likely to avoid punishment [24]. Because the females are smaller, they may have perceived the height to the floor of the rotarod machine as a more salient punishment for quitting, whereas the males were less motivated to remain on the device. A second possible explanation for improved ability to remain on the rotarod could be that the females were significantly smaller and therefore maintained better physical health than their male counterparts. Alhough it is not possible to adequately assess this hypothesis without additional peripheral measures of metabolic health and body composition, our data do support the correlation between physical performance and cognitive outcomes. A PCA revealed poor rotarod performance and weaker grip strength both mapped strongly onto a greater number of trials before reaching criterion on the alternation task.

Previous work utilizing male and female rodents on rotarod-based tasks is scarce, but the limited data available are in alignment with what is demonstrated here. In mice, males performed worse than females at 5 months of age [64]. While male mice continued to decline in rotarod performance with age, female mice did not [37]. In rats, DiFeo and Shors reported that there were no sex differences during puberty, though it appears that early in training females outperformed males [65].These data are essential to investigating age-related cognitive impairments as there is a strong link between frailty and cognitive decline, as well as with longevity [62]. At the very least, our data demonstrate that different parameters are needed for physical assessment across the two sexes, particularly on things in which motivational differences or perceived risk may skew outcomes. Future work should adjust the height of the fall between males and females or include more extensive caloric restriction to manipulate body weight and see the extent to which these parameters interact with sex differences in performance.

Ongoing efforts to assess potential sex differences in behavioral outcomes as well as fundamental investigations into the neurobiological mechanisms of cognitive decline in both males and females are critical to evaluating the efficacy of potential therapeutic interventions. Overall, our data support the idea that there are not fundamental differences in the behavior between females and males on biconditional association and spatial learning and memory tasks, and that both sexes show similar age-related declines. Furthermore, it is not necessary to monitor estrus cycle phase in female rats during behavioral performance of the WM/BAT. That being said, there is an undeniable relationship between estrogen and cognition. In fact, low estrogen is correlated with object recognition impairments that can be restored with estrogen replacement [66,67]. One caveat of the current study is that the aged females were still cycling comparably to the young rats, and may not yet have had the lower levels of estrogen that are associated with female aging. Even with normal cycling, the aged female rats in the current study still demonstrated cognitive decline relative to their young counterparts. This observation indicates that declining ovarian hormones are not the sole mechanism of female cognitive aging, and we need to consider other neurobiological factors of the aging female brain. Because females comprise more than half of the aged population in developed countries such as the United States [9], it is imperative that we include females in aging studies of cognition, and no longer consider the male brain to be the “normal” condition from which females deviate [5].

## Acknowledgements

This research was funded by NIA R01AG049722 (SNB), NIA F31AG058455 (ARH), McKnight Brain Research Foundation, and the University of Florida University Scholars Program (LMT, KTC).

